# mRNA Vaccines Induce Rapid Antibody Responses in Mice

**DOI:** 10.1101/2021.11.01.466863

**Authors:** Makda S. Gebre, Susanne Rauch, Nicole Roth, Janina Gergen, Jingyou Yu, Xiaowen Liu, Andrew C. Cole, Stefan O. Mueller, Benjamin Petsch, Dan H. Barouch

## Abstract

mRNA vaccines can be developed and produced quickly, making them attractive for immediate outbreak responses. Furthermore, clinical trials have demonstrated rapid protection following mRNA vaccination. We sought to investigate how quickly mRNA vaccines elicit antibody responses compared to other vaccine modalities. We first examined immune kinetics of mRNA and DNA vaccines expressing SARS-CoV-2 spike in mice. We observed rapid induction of antigen-specific binding and neutralizing antibodies by day 5 following mRNA, but not DNA, immunization. The mRNA vaccine also induced increased levels of IL-5, IL-6 and MCP-1. We then evaluated immune kinetics of an HIV-1 mRNA vaccine in comparison to DNA, protein, and rhesus adenovirus 52 (RhAd52) vaccines with the same HIV-1 envelope antigen in mice. Induction of envelope-specific antibodies was observed by day 5 following mRNA vaccination, whereas antibodies were detected by day 7-14 following DNA, protein, and RhAd52 vaccination. Eliciting rapid humoral immunity may be an advantageous property of mRNA vaccines for controlling infectious disease outbreaks.

**IMPORTANCE:** mRNA vaccines can be developed and produced in record time. Here we demonstrate induction of rapid antibody responses by mRNA vaccines encoding two different viral antigens by day 5 following immunization in mice. The rapid immune kinetics of mRNA vaccines can be an advantageous property that makes them well suited for rapid control of infectious disease outbreaks.

## INTRODUCTION

In comparison to traditional vaccines, novel mRNA vaccines can be developed and produced for distribution in record time. This makes them attractive candidates for rapidly controlling outbreaks as demonstrated in the current SARS-CoV-2 pandemic [1-4]. Furthermore, clinical trials have now shown the rapid protective efficacy of mRNA vaccines post prime immunizations [5, 6]. For example, the Pfizer mRNA vaccine clinical trial has demonstrated clear divergence between placebo and vaccine recipients only 12 days after the first dose was administered [5].

We sought to investigate how quickly mRNA vaccines induce antibody responses in comparison to other vaccine modalities in mice. Specifically, we immunized C57BL/6 mice intradermally as well as intramuscularly with mRNA or DNA vaccines encoding SARS-CoV-2 full-length pre-fusion stabilized Spike protein [7, 8]. The mRNA vaccine induced binding as well as neutralizing antibody titers as early as 5 days post-immunization. To examine the effect of innate immune triggers, we evaluated the innate cytokine profiles of the two vaccines hours post immunization. Compared to the DNA vaccine, the mRNA vaccine induced a more robust production of IL-5, IL-6 and MCP-1. To determine whether the rapid immune kinetics would translate to other mRNA vaccines of different diseases and antigens, we evaluated the immune kinetics of an mRNA vaccine expressing HIV-1 envelope along with DNA, protein, and Rhesus Adenovirus 52 (RhAd52) vaccines of the same antigen. Again, we were able to observe the rapid induction of antibodies 5 days post mRNA vaccine immunization. The rapid humoral immune kinetics is an advantageous property of the mRNA vaccines, which further supports their use in mitigating infectious disease outbreaks.

## MATERIALS AND METHODS

### Mice and study designs

7-to 8-week-old female C57BL/6 mice (n=5) were purchased from The Jackson Laboratory (Bar Harbor, ME). For SARS-CoV-2 vaccine-based experiments, previously published mRNA (CV2CoV) and DNA vaccines encoding a full-length ancestral SARS-CoV-2 S protein with di-proline mutations were used [8-12]. The mRNA vaccine is obtained from CureVac AG, while the DNA vaccine is produced as previously described [8, 12]. The mRNA vaccine was administered (intramuscularly (I.M.) or intradermally (I.D.)) at 1µg/mouse or 4µg/mouse doses. The DNA vaccine, expressing the same spike antigen, was I.M. injected at 50µg/mouse dose. For the HIV-1 vaccine kinetics experiments, groups of mice were immunized with mRNA (15µg), DNA (50µg), Rhesus Adenovirus 52 (RhAd52) (10^9^ viral particles) or protein vaccines (50µg +100µg Adju phos (InvivoGen)). The HIV mRNA vaccine was also obtained from CureVac AG, while the rest of the vaccines were produced in the Barouch lab. All vaccines encode or represent the same HIV-1 clade C 459C gp140 env antigen. I.D. administrations were administered at 25µL dose at two sites while I.M. injections were administered at 50µL in each of the quadriceps. Blood samples were collected from mice via submandibular bleeds. All animal experiments adhered to the Beth Israel Deaconess Medical Center Institutional Animal Care and Use Committee guidelines.

### ELISA

SARS-CoV-2 Spike as well as HIV-1 Env specific binding antibodies were assessed by Enzyme-linked immunosorbent assays (ELISAs) as described previously [13, 14]. 96-well plates were coated with 1 µg/ml of SARS-CoV-2 S protein (Sino Biological) or HIV-1 clade C Env 459C gp140 in 1× Dulbecco’s phosphate-buffered saline (DPBS) and incubated at 4 °C overnight. The next day, plates were washed once with wash buffer (0.05% Tween-20 in 1× DPBS) and blocked with 350 µl of casein block per well for 2–3 h at room temperature. Next, the block solution was discarded, and plates were blotted dry. Three-fold serial dilutions of serum in casein block were added to wells, and plates were incubated for 1 h at room temperature. Plates were washed three times and then subsequently incubated for 1 h with 0.1 µg ml− 1 of anti-hamster IgG HRP (SouthernBiotech) in casein block at room temperature in the dark. Plates were washed three times, and then 100 µl of SeraCare KPL TMB SureBlue Start solution was added to each well; plate development was halted by the addition of 100 µl of SeraCare KPL TMB Stop solution per well. The absorbance at 450 nm was recorded using a VersaMax or Omega microplate reader. ELISA endpoint titers were defined as the highest reciprocal serum dilution that yielded an absorbance two-fold above background.

### SARS-CoV-2 Pseudovirus neutralization assay

The SARS-CoV-2 pseudoviruses expressing a luciferase reporter gene were generated as described previously [8]. Briefly, the packaging construct psPAX2 (AIDS Resource and Reagent Program), luciferase reporter plasmid pLenti-CMV Puro-Luc (Addgene) and S protein expressing pcDNA3.1-SARS CoV-2 SΔCT were co-transfected into HEK293T cells by lipofectamine 2000 (Thermo Fisher Scientific). The supernatants containing the pseudotype viruses were collected 48 h after transfection; pseudotype viruses were purified by filtration with a 0.45-µm filter. To determine the neutralization activity of the antisera from vaccinated animals, HEK293T-hACE2 cells were seeded in 96-well tissue culture plates at a density of 1.75 × 10^4^ cells per well overnight. Three-fold serial dilutions of heat-inactivated serum samples were prepared and mixed with 50 µl of pseudovirus. The mixture was incubated at 37 °C for 1 h before adding to HEK293T-hACE2 cells. Forty-eight hours after infection, cells were lysed in Steady-Glo Luciferase Assay (Promega) according to the manufacturer’s instructions. SARS-CoV-2 neutralization titers were defined as the sample dilution at which a 50% reduction in relative light units was observed relative to the average of the virus control wells.

### Cytokine Analysis

The levels of 35 cytokines in plasma were determined using U-PLEX Biomarker Group 1 (ms) 35-Plex kit from Meso Scale Discovery (MSD, Rockville, MD). Plasma IFN-α and IFN-β levels were tested using individual U-PLEX Mouse IFN-α Assay and U-PLEX Mouse IFN-β Assay kits from MSD. The lower limit of quantification (LLOQ) of 35 biomarkers are listed as follows: EPO (4.5pg/mL), GM-CSF (0.16 pg/mL), IFN-γ (0.16 pg/mL), IL-1β (3.1 pg/mL), IL-2 (1.1 pg/mL), IL-4 (0.56 pg/mL), IL-5 (0.63 pg/mL), IL-6 (4.8 pg/mL), IL-9 (1.4 pg/mL), IL-10 (3.8 pg/mL), IL-12/IL-23p40 (1.4 pg/mL), IL-12p70 (48 pg/mL), IL-13 (2.7 pg/mL), IL-15 (24 pg/mL), IL-16 (3.6 pg/mL), IL-17A (0.30 pg/mL), IL-17A/F (0.61 pg/mL), IL-17C (2.3 pg/mL), IL-17E/IL-25 (1.6 pg/mL), IL-17F (1.6 pg/mL), IL-21 (6.5 pg/mL), IL-22 (1.2 pg/mL), IL-23 (4.9 pg/mL), IL-27p28/IL-30 (8.7 pg/mL), IL-31 (45 pg/mL), IL-33 (2.2 pg/mL), IP-10 (0.51 pg/mL), KC/GRO (4.8pg/mL) MCP-1 (1.4 pg/mL), MIP-1α (0.21 pg/mL), MIP-1β (13 pg/mL), MIP-2 β (0.30 pg/mL), MIP-3α (0.10 pg/mL), TNF-α (1.3 pg/mL), VEGF-A (0.77 pg/mL), IFN-α (140 pg/mL) and for IFN-β (5.2pg/mL). All samples were run in duplicates. Assays were conducted by Metabolism and Mitochondrial Research Core (Beth Israel Deaconess Medical Center, Boston, MA) following manufacture’s instruction. The assay plates were read by MESO QUICKPLEX SQ 120 instrument and data were analyzed by DISCOVERY WORKBENCH® 4.0 software.

### Statistical analyses

Statistical analyses were performed using GraphPad Prism (version 9.0) software (GraphPad Software) and comparison between groups was performed using a two-tailed nonparametric Mann-Whitney U t test.

## RESULTS

### Rapid induction of binding antibody titers post SARS-CoV-2 Spike mRNA vaccination in mice

To determine the kinetics of humoral immune response, C57BL/6 mice were vaccinated intramuscularly (I.M.) or intradermally (I.D.) with mRNA vaccines expressing SARS-CoV-2 Spike at doses of 1 µg or 4 µg. Additional groups of mice were immunized I.M. with a previously investigated DNA vaccine [8] at a dose of 50µg or with PBS as a sham control. Spike-specific binding antibodies were measured in serum by ELISA. Spike-specific binding antibody titers (median 179; range 72 – 532) were observed by day 5 following I.D. immunization with the 4µg dose of the mRNA vaccine (Figure 1). Antibody titers were also observed 5 days post I.M. immunization, although I.D. immunization resulted in higher titers at this early timepoint (P = 0.0317). In contrast, the median titers from both low dose mRNA immunization and DNA vaccination remain below limit of detection on day 5. By day 7, we still observed significantly higher antibody titers for mice immunized I.M. with the 4µg dose of mRNA compared to mice immunized I.M. with the DNA vaccine (P=0.0079). On day 14, all mRNA groups elicited dose-dependent antibody responses with no significant difference between I.M. and I.D. vaccination. Lower but detectable titers were also observed in the DNA vaccine group, while the sham group antibody titers remained at baseline. By day 21, antibody titers elicited by the DNA vaccine were comparable to those induced by the 4µg dose mRNA groups. These results demonstrate that mRNA vaccines induce rapid antibody responses and that these responses are dose and route dependent.

**Figure 1.**
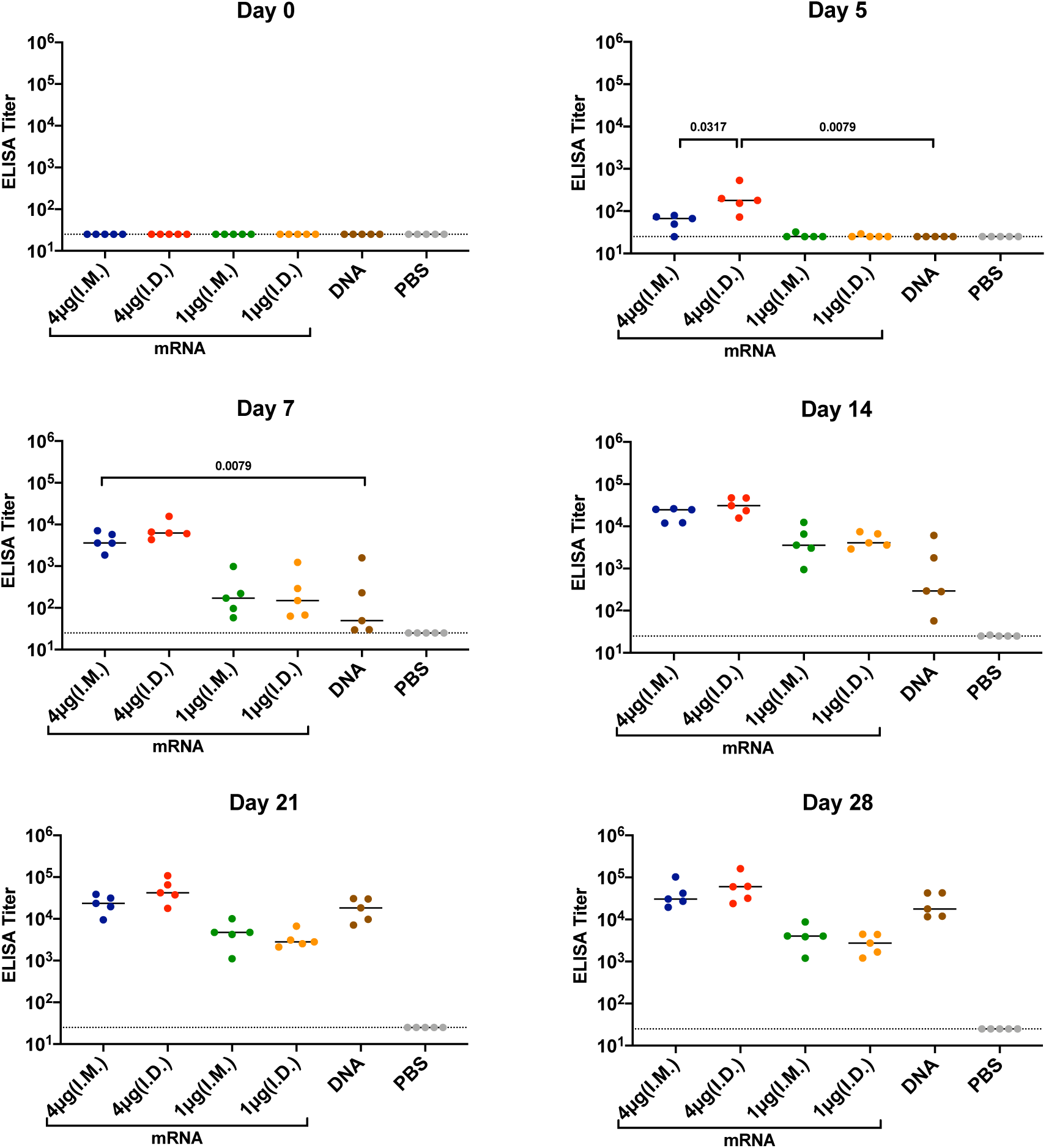
Kinetics of binding antibody responses of mice immunized with SARS-CoV-2 Spike mRNA and DNA vaccines. C57BL/6 mice were immunized I.M. or I.D. with spike encoding mRNA vaccine (1µg or 4µg/mouse), DNA vaccine (50µg/mouse) or PBS. Binding antibody titers were assessed via ELISA at 0, 5, 7, 14, 21 and 28 days post immunization. Each dot represents an individual animal, bars depict the median and the dotted line shows limit of detection. Statistical analysis was performed using Mann-Whitney test. (I.M = intramuscular; I.D. = intradermal)

### Early Induction of neutralizing antibodies upon SARS-CoV-2 Spike mRNA vaccination

Next we evaluated neutralizing antibody titers via a pseudovirus neutralization assay [15]. Consistent with the binding antibody titer data, mice immunized via either I.M. or I.D. routes at the 4µg dose of mRNA vaccine exhibited neutralizing antibody (NAb) titers as early as day 5 following immunization (Figure 2). Mice immunized I.D. with the 4µg dose exhibited the highest NAb titers on day 5 (median 79; range 24 – 221). By day 7, mice immunized I.M. with the 4µg of mRNA vaccine showed significantly higher neutralizing antibody titers than mice immunized I.M. with the DNA vaccine (P=0.0079). The 1µg dose mRNA vaccine elicited lower NAbs than the 4µg dose. By day 21, NAb titers induced by the DNA vaccine reached levels that were comparable to those elicited by the 4µg mRNA vaccine.

**Figure 2.**
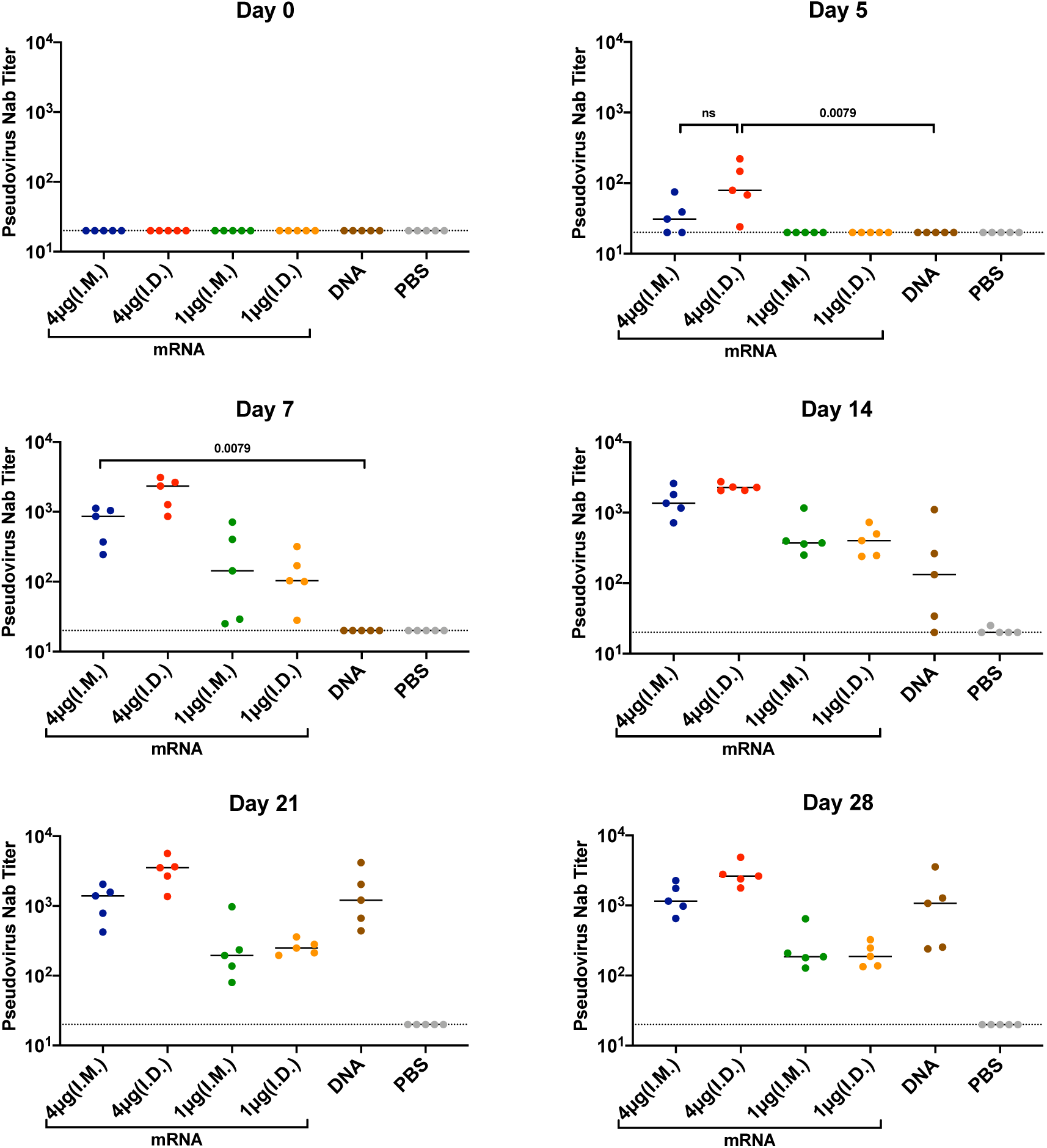
Kinetics of neutralizing antibody titers of mice immunized with SARS-CoV-2 Spike mRNA and DNA vaccines. C57BL/6 mice were immunized I.M. or I.D. with spike encoding mRNA vaccine at 1µg/mouse or 4µg/mouse doses, spike encoding DNA at 50µg/mouse dose or PBS. Neutralizing antibody titers were assessed via pseudovirus neutralization assay at 0, 5, 7, 14, 21 and 28 days post immunization. Each dot represents an individual animal, bars depict the median and the dotted line shows limit of detection. Statistical analysis was performed using Mann-Whitney test. (I.M = intramuscular; I.D. = intradermal)

### Cytokine and Chemokine responses of mRNA and DNA vaccines

To better understand differences in innate immune responses post SARS-CoV-2 mRNA or DNA vaccination that may contribute to humoral immune responses, we evaluated the kinetics of expression of cytokines and chemokines at 5 and 24 hours post vaccination in mouse plasma samples. We compared mice that were immunized with 4µg of mRNA vs 50µg of DNA via I.M. injection routes. Cytokines that play a key role in the initiation and development of B cells and antibody production were significantly induced following mRNA vaccination (Figure 3). IL-5, which is a critical cytokine for mouse B cell differentiation to antibody-secreting plasma cells [16], was higher (P=0.0317) in mRNA vaccinated mice (median 14.62pg/mL) than in DNA vaccinated mice (median 6.41pg/mL) at 5 hours following immunization (Figure 3). Similarly, IL-6, which is critical for B cell proliferation and isotype switching, was higher (P=0.0079) 5 hours following mRNA vaccination (median 92.37pg/mL) compared to DNA vaccination (median 12.43pg/mL) [17]. Furthermore, MCP-1, MIP-1α, MIP-1β as well as IP-10, which are key chemokines for antigen presenting cell activation and migration, were also induced at 5 and 24 hours after mRNA immunization.

**Figure 3.**
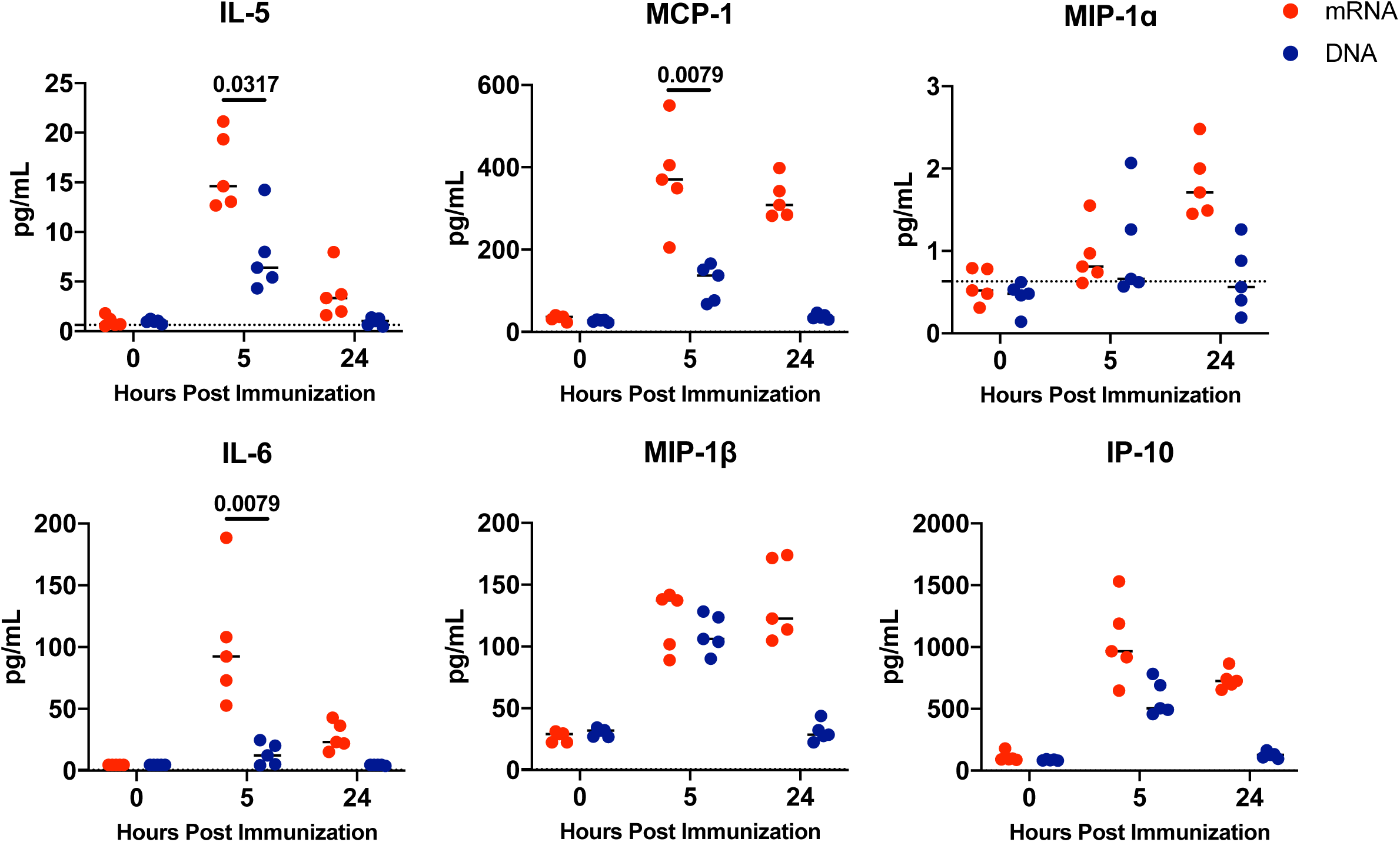
Cytokine and chemokine responses elicited in mice immunized with SARS-CoV-2 Spike mRNA and DNA vaccines. C57BL/6 mice were immunized I.M. with either 4µg/mouse dose of spike encoding mRNA vaccine or, 50µg/mouse dose of spike encoding DNA vaccine. Plasma collected at 0, 5 and 24 hours post immunization were analyzed for cytokines using a U-PLEX Biomarker Group 1 (ms) 35-Plex kit from Meso Scale Discovery. Each dot represents an individual animal, bars depict the median and the dotted line shows limit of detection. Statistical analysis was performed using Mann-Whitney test. (I.M = intramuscular)

### Humoral immune kinetics in mice immunized with HIV-1 env mRNA vaccine

We next sought to determine whether the rapid kinetics observed with SARS-CoV-2 Spike mRNA vaccine could be generalizable to mRNA vaccines encoding other antigens. We evaluated the antibody kinetics of an mRNA vaccine (15µg/mouse) encoding a Clade C 459C HIV-1 envelope (Env) gp140, as well as a DNA vaccine (50µg/mouse), purified protein vaccine (50µg/mouse with 100µg Adjuphos adjuvant), and rhesus adenovirus 52 (RhAd52) vaccine (10^9^ viral particles/mouse) with the same HIV Env antigen. These vaccines were all delivered I.M. Low antibody titers (median 52; range 32 – 66) were observed on day 3 following HIV-1 Env mRNA immunization, but robust antibody responses (median 356; range 114 – 689) were evident on day 5 following HIV-1 mRNA vaccine immunization, whereas DNA, protein, and RhAd52 vaccines were largely baseline at that time point (P=0.0079) (Figure 4). By day 21, all four vaccine modalities showed similar robust antibody titers. These data suggest that immune response kinetics after mRNA vaccination are more rapid than with three other leading vaccine technologies.

**Figure 4.**
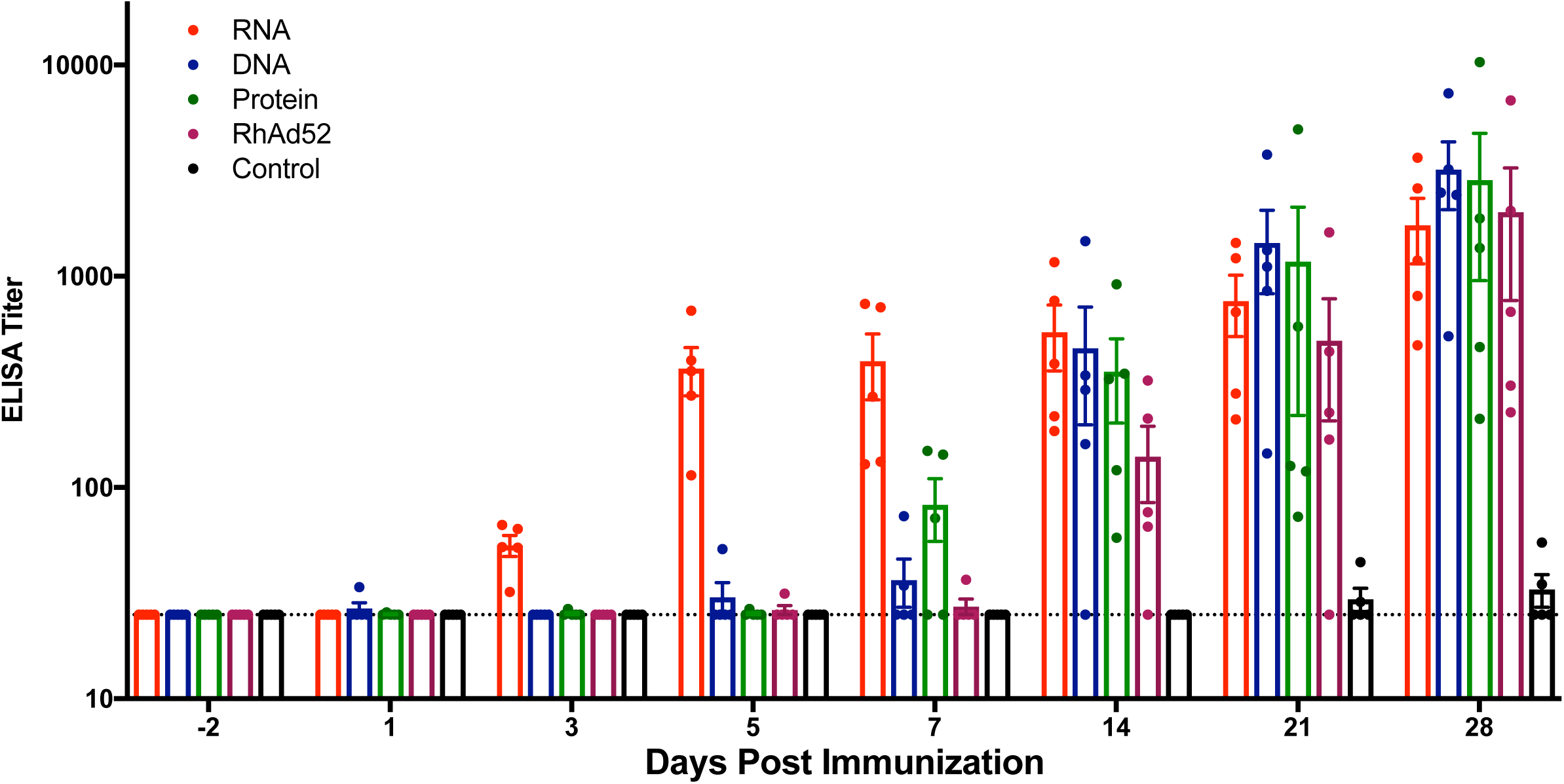
Rapid humoral immunity of mRNA vaccine expressing gp140WT compared to DNA, RhAd52 and protein vaccine modalities of the same antigen. C57BL/6 mice were immunized I.M. with mRNA (15µg), DNA (50µg), rhesus adenovirus 52 (RhAd52) (10^9^ viral particles) or protein (50µg +100µg Adju-phos (InvivoGen) vaccines that encode or represent the HIV-1 env antigen. Antigen specific binding antibodies were assessed at -2, 1, 3, 5, 7, 14, 21, and 28 days post immunization via ELISA. Each dot represents an individual animal, bars depict the median and the dotted line shows limit of detection. Statistical analysis was performed using Mann-Whitney test. (I.M = intramuscular)

## DISCUSSION

Novel mRNA vaccines have demonstrated remarkable utility in controlling infectious disease outbreaks. They can be developed and produced at a large scale more rapidly than other traditional vaccine methods. In this study, we assessed the kinetics of induction of antibody responses following immunization of mice with mRNA vaccines expressing SARS-CoV-2 spike or HIV-1 envelope. Both mRNA vaccines elicited rapid antibody responses by day 5 following immunization.

Multiple studies have demonstrated that mRNA vaccines elicit strong antibody responses in various mouse models [18-21], as well as robust germinal center responses [22] that may be associated with neutralizing antibody generation [23]. Although a wealth of information has been gained regarding antibody responses induced by mRNA vaccines at later timepoints, the early and immediate humoral immune kinetics of mRNA vaccines remains unclear. We detected antigen specific antibodies by day 5 after immunization in mice following SARS-CoV-2 spike mRNA vaccination but not DNA immunization. Neutralizing antibodies were also detected at this early time point. The early antibody responses were both dose and route dependent. Intradermal immunization of the mRNA vaccine elicited higher early antibody responses than intramuscular immunization at the 4µg dose, which is consistent with prior studies that have reported that I.D. immunization of mRNA vaccines results in stronger antibody responses than I.M. immunization [24, 25]. This may be due to local antigen-presenting cells (APCs) in the skin, such as dermal dendritic cells, to process and deliver antigen to T and B cells in the draining lymph nodes [26]. In Thailand, I.D. administration of the rabies vaccine for post-exposure prophylaxis (PEP) has offered a cost-effective alternative to I.M. immunizations [27].

To examine differences in early innate responses induced by mRNA and DNA vaccines, we compared cytokine profiles at 5 and 24 hours post immunization. Cytokines that play a key role in B cell development, such as IL-5 and IL-6, were better induced following mRNA vaccination. IL-5 supports terminal mouse B cell differentiation to antibody-secreting plasma cells and promotes homeostatic proliferation, survival and antibody production [16]. IL-6 is critical for B cell proliferation and isotype switching, which is necessary to produce IgG antibodies that are represented in our antibody titer data [17]. Other cytokines that are upregulated post mRNA vaccine immunization, such as MCP-1, MIP-1α, MIP-1β as well as IP-10, are important for APC recruitment and activation. MCP-1 recruits monocytes, memory T cells, and dendritic cells to the sites of inflammation [28]. MIP-1α and MIP-1β are major cytokines produced by macrophages and monocytes during inflammation and promote lymphocyte migration [29, 30]. IP-10 is attributed to several roles, such as chemoattraction for monocytes/macrophages, T cells, NK cells, and dendritic cells, promotion of T cell adhesion to endothelial cells [31]. Overall, the cytokine data suggests that mRNA vaccination induces cytokines that are key in APC recruitment and activation as well as B cell differentiation and proliferation.

We show that rapid antibody responses were induced not only by mRNA vaccines for SARS-CoV-2 spike but also for HIV envelope, suggesting the generalizability of our findings. mRNA vaccines for HIV have previously been investigated, although the kinetics of early antibody production has not been examined [32]. Our results suggest that rapid humoral response is a characteristic of the mRNA vaccine platform rather than an antigen-specific finding. Other vaccine platforms, such as DNA, RhAd52, and protein vaccines, are also immunogenic but do not show the rapid induction of antibody responses by day 5 following immunization.

In summary, our data show that two different mRNA vaccines induced rapid antibody responses by day 5 following immunization in mice. These findings may help explain the rapid protection achieved with mRNA vaccines in clinical trials, which is useful in containing infectious isease outbreaks.

## ACKNOWLEDGMENTS

We thank J. Ventura and L. Tostanoski for generous advice and assistance. We acknowledge funding from the National Institute of Health (AI124377, AI128751, AI126603, CA260476), CureVac, and the Ragon Institute of MGH, MIT, and Harvard.

## REFERENCES

1. Cucinotta, D. and and. Vanelli, WHO Declares COVID-19 a Pandemic. Acta Biomed, 2020. 91(1): p. 157–160.

2. Pardi, N., et al., mRNA vaccines - a new era in vaccinology. Nat Rev Drug Discov, 2018. 17(4): p. 261–279.

3. Rauch, S., et al., New Vaccine Technologies to Combat Outbreak Situations. Front Immunol, 2018. 9: p. 1963.

4. Gebre, M.S., et al., Novel approaches for vaccine development. Cell, 2021. 184(6): p. 1589–1603.

5. Polack, F.P., et al., Safety and Efficacy of the BNT162b2 mRNA Covid-19 Vaccine. N Engl J Med, 2020. 383(27): p. 2603–2615.

6. Baden, L.R., et al., Efficacy and Safety of the mRNA-1273 SARS-CoV-2 Vaccine. N Engl J Med, 2021. 384(5): p. 403–416.

7. Hoffmann, D., et al., CVnCoV and CV2CoV protect human ACE2 transgenic mice from ancestral B BavPat1 and emerging B.1.351 SARS-CoV-2. Nat Commun, 2021. 12(1): p. 4048.

8. Yu, J., et al., DNA vaccine protection against SARS-CoV-2 in rhesus macaques. Science, 2020. 369(6505): p. 806–811.

9. Kirchdoerfer, R.N., et al., Pre-fusion structure of a human coronavirus spike protein. Nature, 2016. 531(7592): p. 118–21.

10. Wrapp, D., et al., Cryo-EM structure of the 2019-nCoV spike in the prefusion conformation. Science, 2020. 367(6483): p. 1260–1263.

11. Rauch, S., et al., mRNA-based SARS-CoV-2 vaccine candidate CVnCoV induces high levels of virus-neutralising antibodies and mediates protection in rodents. NPJ Vaccines, 2021. 6(1): p. 57.

12. Nicole Roth, J.S., Donata Hoffmann, Moritz Thran, Andreas Thess, Stefan Stefan. Mueller, Benjamin Petsch and Susanne Rauch, CV2CoV, an enhanced mRNA-based SARS-CoV-2 vaccine candidate, supports higher protein expression and improved immunogenicity in rats. bioRxiv, 2021.

13. Tostanoski, L.H., et al., Ad26 vaccine protects against SARS-CoV-2 severe clinical disease in hamsters. Nat Med, 2020. 26(11): p. 1694–1700.

14. Nkolola, J.P., et al., Breadth of neutralizing antibodies elicited by stable, homogeneous clade A and clade C HIV-1 gp140 envelope trimers in guinea pigs. J Virol, 2010. 84(7): p. 3270–9.

15. Yu, J., et al., Deletion of the SARS-CoV-2 Spike Cytoplasmic Tail Increases Infectivity in Pseudovirus Neutralization Assays. J Virol, 2021.

16. Takatsu, K., T. Kouro, and and. Nagai, Interleukin 5 in the link between the innate and acquired immune response. Adv Immunol, 2009. 101: p. 191–236.

17. Vazquez, M.I., J. Catalan-Dibene, and and. Zlotnik, B cells responses and cytokine production are regulated by their immune microenvironment. Cytokine, 2015. 74(2): p. 318–26.

18. Ji, R.R., et al., BNT162b2 Vaccine Encoding the SARS-CoV-2 P2 S Protects Transgenic hACE2 Mice against COVID-19. Vaccines (Basel), 2021. 9(4).

19. Huang, Q., et al., A single-dose mRNA vaccine provides a long-term protection for hACE2 transgenic mice from SARS-CoV-2. Nat Commun, 2021. 12(1): p. 776.

20. Corbett, K.S., et al., SARS-CoV-2 mRNA vaccine design enabled by prototype pathogen preparedness. Nature, 2020. 586(7830): p. 567–571.

21. Vogel, A.B., et al., BNT162b vaccines protect rhesus macaques from SARS-CoV-2. Nature, 2021. 592(7853): p. 283–289.

22. Pardi, N., et al., Nucleoside-modified mRNA vaccines induce potent T follicular helper and germinal center B cell responses. J Exp Med, 2018. 215(6): p. 1571–1588.

23. Lederer, K., et al., SARS-CoV-2 mRNA Vaccines Foster Potent Antigen-Specific Germinal Center Responses Associated with Neutralizing Antibody Generation. Immunity, 2020. 53(6): p. 1281–1295 e5.

24. Pardi, N., et al., Expression kinetics of nucleoside-modified mRNA delivered in lipid nanoparticles to mice by various routes. J Control Release, 2015. 217: p. 345–51.

25. Bahl, K., et al., Preclinical and Clinical Demonstration of Immunogenicity by mRNA Vaccines against H10N8 and H7N9 Influenza Viruses. Mol Ther, 2017. 25(6): p. 1316–1327.

26. Zeng, C., et al., Formulation and Delivery Technologies for mRNA Vaccines. Curr Top Microbiol Immunol, 2020.

27. Gongal, G. and and. Sampath, Introduction of intradermal rabies vaccination - A paradigm shift in improving post-exposure prophylaxis in Asia. Vaccine, 2019. 37 Suppl 1: p. A94–A98.

28. Deshmane, S.L., et al., Monocyte chemoattractant protein-1 (MCP-1): an overview. J Interferon Cytokine Res, 2009. 29(6): p. 313–26.

29. Schaller, T.H., et al., Chemokines as adjuvants for immunotherapy: implications for immune activation with CCL3. Expert Rev Clin Immunol, 2017. 13(11): p. 1049–1060.

30. Lillard, J.W., Jr., et al., MIP-1alpha and MIP-1beta differentially mediate mucosal and systemic adaptive immunity. Blood, 2003. 101(3): p. 807–14.

31. Costela-Ruiz, V.J., et al., SARS-CoV-2 infection: The role of cytokines in COVID-19 disease. Cytokine Growth Factor Rev, 2020. 54: p. 62–75.

32. Mu, Z., B.F. Haynes, and D.W. Cain, HIV mRNA Vaccines-Progress and Future Paths. Vaccines (Basel), 2021. 9(2).

